# A biophysical model uncovers the size distribution of migrating cell clusters across cancer types

**DOI:** 10.1101/563049

**Authors:** Federico Bocci, Mohit Kumar Jolly, José Nelson Onuchic

## Abstract

Migration from the primary tumor is a crucial step in the metastatic cascade. Cells with various degrees of adhesion and motility migrate and are launched into the bloodstream as single circulating tumor cells (CTCs) or multi-cellular CTC clusters. The frequency and size distributions of these clusters has been recently measured, but the underlying mechanisms enabling these different modes of migration remain poorly understood. We present a biophysical model that couples the phenotypic plasticity enabled by the epithelial-mesenchymal transition (EMT) and cell migration to explain the modes of individual and collective cancer cell migration. This reduced physical model undergoes a transition from individual migration to collective cell migration and robustly recapitulates CTC cluster fractions and size distributions observed experimentally across several cancer types, thus suggesting the existence of common features in the mechanisms underlying cancer cell migration. Furthermore, we identify mechanisms that can maximize the fraction of CTC clusters in circulation. First, mechanisms that prevent a complete EMT and instead increase the population of hybrid Epithelial/Mesenchymal (E/M) cells are required to recapitulate CTC size distributions with large clusters of 5-10 cells. Moreover, multiple intermediate E/M states give rise to larger and heterogeneous clusters formed by cells with different epithelial-mesenchymal traits. Overall, this biophysical model provides a platform to continue to bridge the gap between the molecular and biophysical regulation of cancer cell migration, and highlights that a complete EMT might not be required for metastasis.

## Introduction

Metastases are the leading cause of cancer-related deaths and still represent an insuperable clinical challenge [1]. Typically, the epithelial cells in a primary tumor gain some degree of motility that enables migration through the adjacent tissues, invasion to breach the basement membrane and enter the bloodstream, and eventually colonize a distant organ to start a metastasis. Individually migrating cells - or circulating tumor cells (CTCs) - have been considered the primary drivers of metastasis. Recently, however, experimental evidence suggests that cells can also plunge into the bloodstream as multi-cellular clusters, which can more efficiently survive through the metastatic cascade and initiate a secondary tumor [2,3]. In animal models, these clusters have been shown to initiate more than 95% of all metastases, despite accounting for as low as 3% of all CTC events (individual and clusters taken together) and typically only 5-8 cells large [2,3].

The epithelial-mesenchymal transition (EMT) is among the most important mechanisms that confers motility to epithelial cancer cells and enables migration [4,5]. During EMT, cells partially or completely lose E-cadherin-mediated adhesion junctions and apicobasal polarization while gaining motility and invasive traits [4]. Moreover, cancer cells can display a spectrum of EMT states [6], including many hybrid epithelial/mesenchymal (E/M) cell states [7]. These hybrid states can conserve cell-cell adhesion typical of epithelial cells while acquiring the motility typical of mesenchymal cells, hence allowing collective migration leading to the formation of clusters of CTCs [6]. So far, however, no quantitative model has been proposed to explain how a partial or complete EMT could potentially translate into the different frequencies and size distributions of CTC clusters observed in different tumors.

Here, we devise a biophysical model to investigate various modes of cancer cell migration emerging from different degrees of phenotypic plasticity enabled by a partial or complete EMT. In our model, cancer cells can undergo a complete EMT which only allows solitary migration, but can also attain an intermediate phenotype that enables collective migration. These premises can explain the individual and clustered cell migration observed in different cancer types and robustly recapitulate the fraction and size distribution of CTC clusters experimentally observed in patients and mouse models, therefore offering a common framework for quantifying cell migration across different cancer types.

Furthermore, given the connection between hybrid epithelial/mesenchymal cell state(s), CTC clusters and poor prognosis [8], we seek mechanisms that increase the fraction of CTC clusters and allow the formation of larger clusters. First, a partial EMT facilitates the formation of large clusters and quantitatively reproduces CTC cluster distributions with large clusters that are not captured by a model of complete EMT. Therefore, our model suggests that a partial EMT characterized by conservation of an epithelial program might be sufficient to explain invasion in multiple cancers. Moreover, multiple intermediate hybrid E/M states facilitate the formation of large clusters and promote cell heterogeneity, mechanisms that can lead to higher plasticity, aggressiveness, and drug resistance [6].

## Methods - Building a biophysical model for cancer invasion

In order to metastasize, cancer cells undergo a sequence of genetic, epigenetic, and physical transformations. First, epithelial cells in a tumor must transition to a phenotype enabling some degree of motility. This phenotype, in concerto with interactions with the extracellular matrix (ECM), modulates a solitary or collective migration. Afterwards, these cells intravasate to access the circulatory system, where they can face immune recognition and external drugs. We build a biophysical model that can integrate different aspects of cancer cell migration at a phenomenological level with parameters that are related to biological processes.

To model phenotypic plasticity within a tumor, we consider a scenario where cells in a cell layer can assume three possible states. First, an epithelial cell state (E) that has maximal adhesion to neighboring cells and does not allow migration. Second, an intermediate cell state, that we will refer to as a Hybrid epithelial/mesenchymal (E/M or H), characterized by cell adhesion and migration. Third, a fully differentiated mesenchymal (M) cell state that has no adhesion with neighbors and can only migrate solitarily. Cells can switch from E to E/M state in partial EMT transitions and from E/M to M state in complete EMT transitions with rate k (Fig. 1A). The model can be generalized to more than one intermediate state.

**Figure 1.**
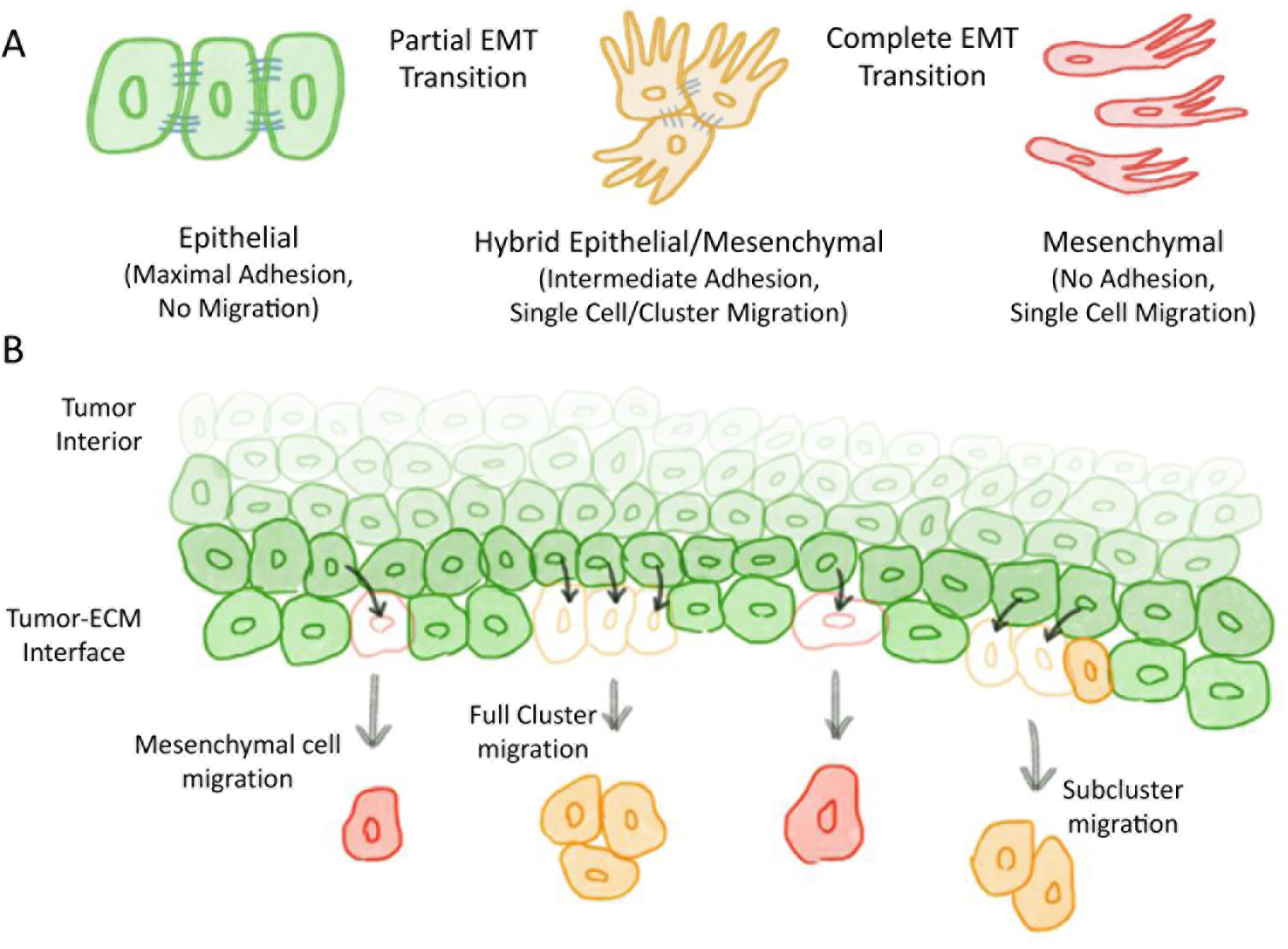
A biophysical model to couple EMT phenotype transitions and cell migration. **(A)** Cell phenotype transitions in the model. **(B)** Migration and replacement of new epithelial cells in the model. When two or more hybrid E/M cells are neighbors, the entire cluster can migrate together (full cluster escape) or can break into smaller subcluster(s) that migrate(s) alone (subcluster escape).

Second, to model cell migration, E/M and M cells can migrate from the cell layer (Fig. 1B). M cells can only migrate as single cells due to complete loss of E-cadherin-based cell adhesion. Conversely, two or more E/M cells can also migrate as a multi-cellular cluster if they are in spatial proximity, due to partial conservation of their adhesion program [9]. In order to migrate, cells need to break the adhesive bonds with their epithelial (E) and/or hybrid E/M neighbors. Unbinding between cells is modeled as a stochastic event that take place with probabilities p_E_ (breaking of an E-H bond) and p_H_ (breaking of an H-H bond) (Supplementary Information A). As hybrid E/M and mesenchymal cells migrate from the lattice, they are replaced by new epithelial cells. It is assumed that cells in the tumor interior are ‘shielded’ from EMT-inducing signals, and can undergo EMT only once they reach the periphery of the tumor [10,11].

Furthermore, a cooperativity index (c) regulates how preferable it is for the E/M cells in a cluster to migrate together (Supplementary Information A). In the limit case where c=0, there is no preference for collective migration. The case of c=1 implies a purely additive motility (i.e. a cluster of 2 E/M cells has twice the motility of a single E/M cell), while c>1 implies cooperation between E/M cells during migration (i.e. the motility of a 2-cell cluster is more than twice the motility of a single cell). This term integrates different processes. First, hybrid E/M cells that share adhesive bonds can carry each other during migration, hence making collective migration more likely. Furthermore, the stiffness and spatial organization of the surrounding ECM can select a more solitary or collective migration [12,13].

Therefore, the probability to migrate for clusters of various sizes is determined by different factors. First, the probability that such clusters exist in the cell layer, which depends on the size of the hybrid E/M cell population. Second, the probability to break the adhesive bonds with the neighboring cells. Lastly, the strength of collective pushing and modulation by the ECM (Supplementary Information B).

We do not explicitly consider the effect of intravasation. Although intravasation is likely to act as a ‘filter’ for migrating CTCs, the current work is mostly concerned with small clusters of up to 5-10 cells. *In vitro* experiments show that small clusters can effectively rearrange their conformation to traverse capillary-sized vessels [14]. Similarly, we do not explicitly include cell death in circulation. It is assumed that this term depends on the cell phenotype through antigen recognition at cell surface, rather than on the size of the CTC clusters, and therefore reduces the total number of CTCs but does not modify the relative frequency of CTCs of different sizes.

Although tumors have a three-dimensional structure, the periphery of a tumor where cells are free to migrate can be modeled as a two-dimensional surface. However, the probability to observe clusters of various sizes in a two-dimensional cell layer cannot be calculated analytically. For this reason, we implement an ‘effective-2D’ approximation that uses the one-dimensional analytical calculation of the cluster size probability distribution but (i) makes the model more realistic by accounting for the larger number of nearest neighbors (and, therefore, adhesive contacts to break) in a two-dimensional lattice and (ii) restricts the computational cost of a simulation on a two-dimensional lattice (Supplementary Information C-E and Fig. S1). The code developed to study the model is freely available at https://github.com/federicobocci91.

## Results

### A transition from single cell migration to clustered cell migration

First, we characterize the behavior of the model in terms of population of Epithelial (E), Hybrid E/M, and Mesenchymal (M) cells by varying the rate of EMT *k*. This dimensionless parameter represents how slow or fast EMT is as compared to the typical timescale required for cell migration; we will simply refer to k as the EMT rate to simplify the notation (Supplementary Information F). In presence of a slow EMT (*k* ≪ 1), most cells are epithelial, while a fast EMT (*k* ≫ 1) converts most cells to a mesenchymal state (Fig. 2A). However, an intermediate EMT rate (*k* ≈ 1) allows a high fraction of E/M cells (Fig. 2A), and therefore makes the contact between E/M cells and the formation of clusters more probable. To quantify this concept, we define a ‘cluster escape fraction’ as the fraction of migration events from the cell layer that involve a cluster rather than a single cell. As expected, this quantity is maximized at intermediate levels of the EMT rate (Fig. 2B).

**Figure 2.**
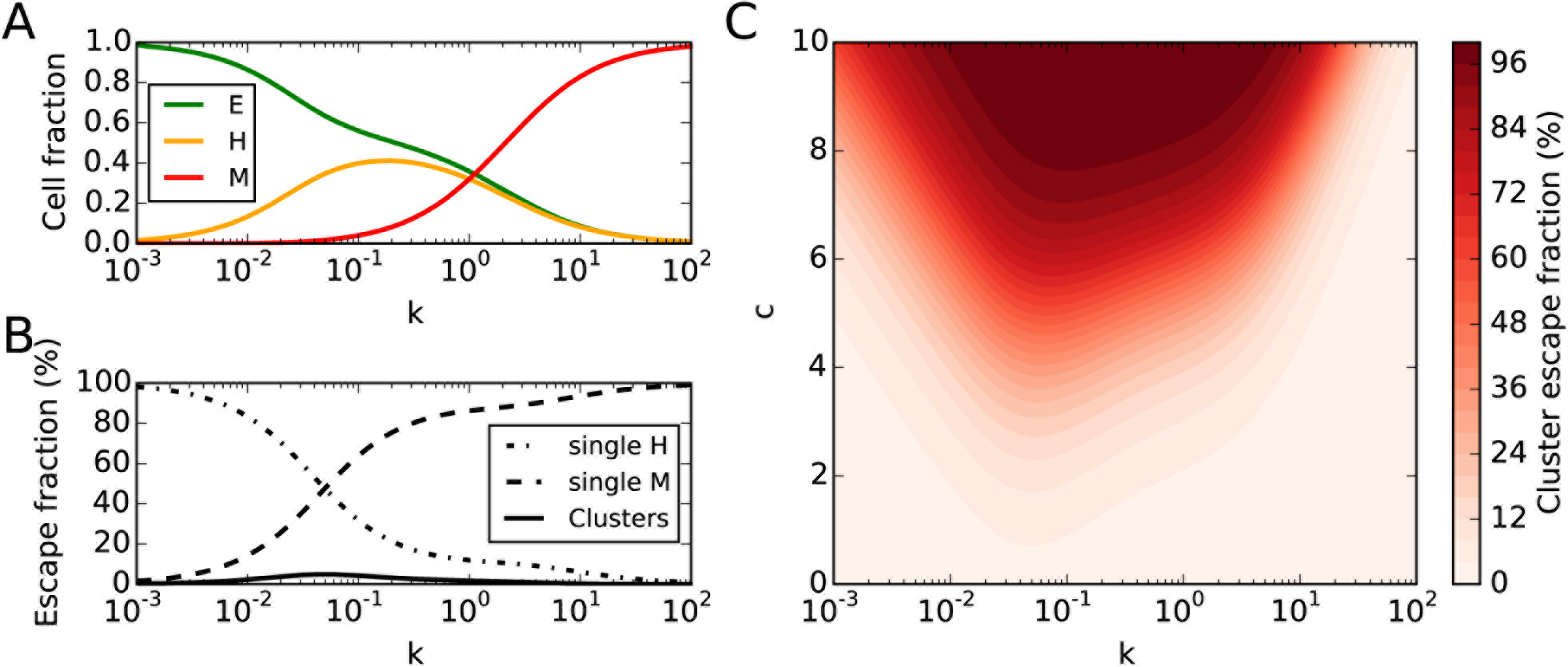
EMT rate and migration cooperativity give rise to a switch from single cell migration to cluster migration. **(A)** Fraction of epithelial (E, green), hybrid E/M (H, yellow) and mesenchymal (M, red) cells as a function of the rate of EMT (k). **(B)** Fractions of escape events due to E/M clusters, single E/M (H) cells and single M cells as a function of k. **(C)** Cluster escape fraction as a function of k (x-axis) and the migration cooperativity index (c, y-axis). c=1 in panels A, B.

A more systematic analysis of the model’s behavior for varying combinations of parameters highlights a switch from single cell migration to clustered cell migration. Specifically, the cluster escape fraction increases as a combination of two factors. First, when the rate of EMT (k) is neither too high nor too low, ensuring a high fraction of hybrid E/M cells in the lattice; second, a high migration cooperativity (c) that promotes collective, rather than solitary, cell migration (Fig. 2C and S2A). These conditions ensure not only a larger cluster escape fraction, but also increase the typical cluster size to up to 5 cells (Fig. S2B). These trends are robust to variation of the model’s parameters and contact geometry between cells (Fig. S2C-D, S3).

This variability in the frequency of CTC clusters is well-supported by multiple independent experiments that reported variable fractions of CTC clusters in circulation [2,15–19]. These CTC cluster fractions indicate that different cancer types are characterized by different migratory regimes, that in turn correspond to different combinations of EMT rate and cooperativity (k, c) in the model (Fig. S4). Potentially, different cancer subtypes or even patient-specific factors might further increase this variability.

### The difference between maximal CTC cluster fraction and maximal CTC count

The fraction of escape events that involve clusters or single cells is not directly informative on the absolute ability of a tumor to disseminate CTCs, that is quantified experimentally by the total count of CTCs per unit volume of blood [20]. The rate of escape of clusters is maximal at intermediate levels of the EMT rate (k), i.e. the regime where the population of hybrid E/M cells is larger (Fig. 3A). However, the total migration rate, which considers single E/M cells, E/M clusters and single mesenchymal cell, is maximal at a much higher EMT rate (Fig. 3B). When the rate of EMT is very fast, most cells acquire a mesenchymal state and thus tend to migrate as single cells.

**Figure 3.**
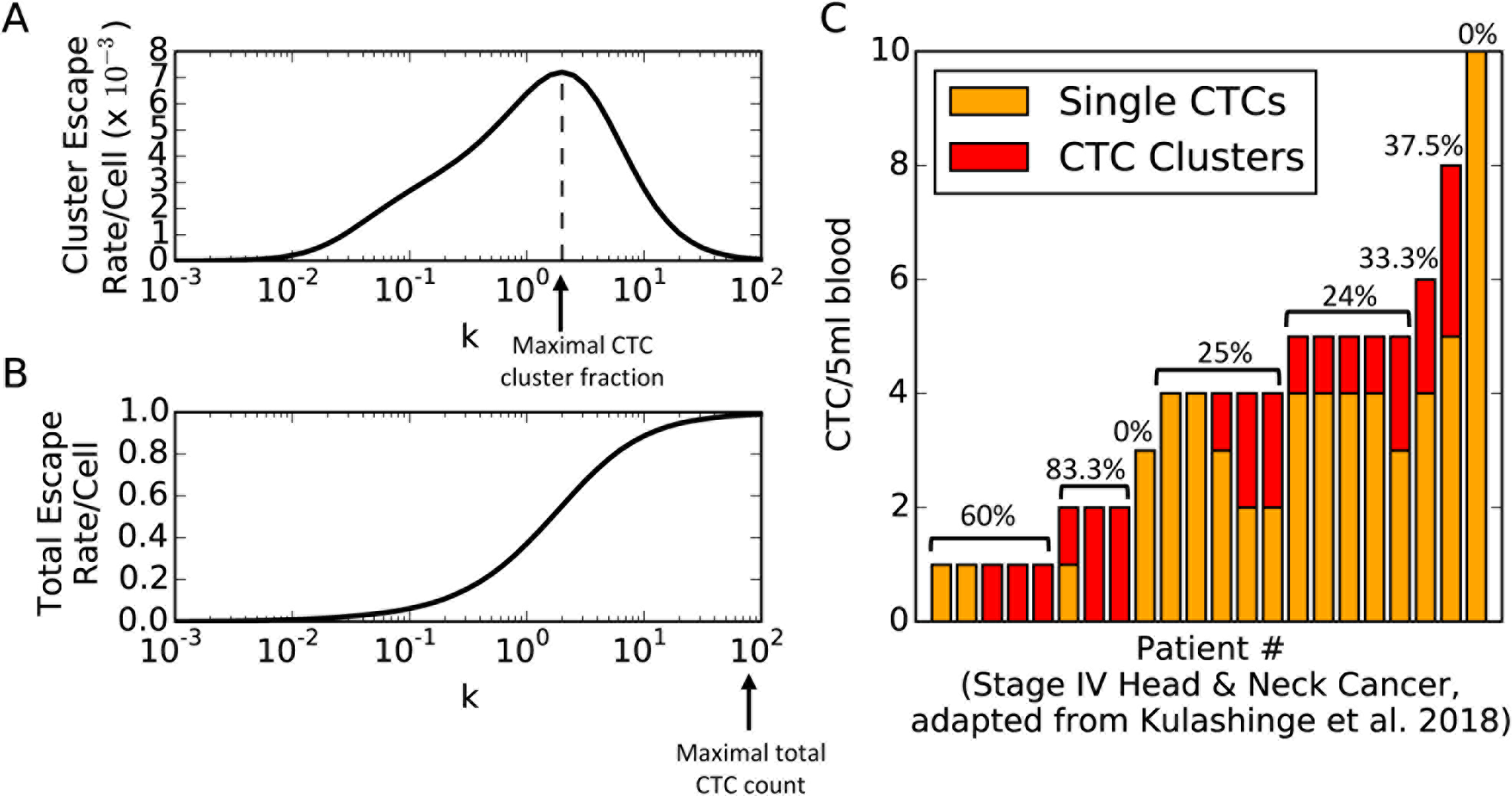
Separation between the regimes of maximal cluster dissemination and maximal overall dissemination. **(A)** Cluster escape rate and **(B)** total escape rate (clusters + single cells) per unit cell as a function of the EMT rate (k). **(C)** Number of single CTCs and CTC clusters measured in circulation of Stage IV head and neck cancer patients. Patients are ranked based on the total number of CTCs detected. Percentages indicate CTC cluster fraction for groups with same total CTC count.

The separation between these two regimes can be appreciated in a clinical dataset that quantifies the number of single CTCs and CTC clusters in the bloodstream of stage IV head and neck cancer patients [19]. When ranking patients based on their total CTC number, it is evident that most patients with lower total CTC count have a higher percentage of CTC clusters (Fig. 3C).

The metastatic potential of a tumor is likely to be a trade-off between slower dissemination of highly-metastatic but somewhat rarer hybrid E/M clusters and faster dissemination of less metastatic individually migrating mesenchymal cells. Our model predicts a separation between regimes with high CTC cluster fraction and regimes with high total CTC count. To verify this prediction and, more importantly, develop reliable prognostic tools, future investigations will require robust measurement of the CTC absolute number as well as fraction and size distribution of CTC clusters.

### The single cell-cluster migration transition recapitulates cluster size distributions across several cancer types

So far, we have examined the overall fraction of migrating clusters and compared it to experimental measurements on CTC cluster fraction and counting. Experimental techniques, however, have recently gained the sensibility to measure the size distribution of CTC clusters in various cancers, highlighting a tremendous variability that can be explored through the lens of our biophysical model (Supplementary Information G).

First, data from melanoma patients acquired via microfluidic trap [21] shows a very steep distribution that is reproduced well by a model with low cooperativity (c=3) (Fig. 4A). Similarly, a distribution from prostate cancer [15] shares similar traits but slightly larger tail, and maps well on to a larger cooperativity index (c=4) (Fig. 4B). Microfluidic data of breast cancer patients [21] shows a more prominent role of large clusters, where approximately two-thirds of the observed clusters are composed by more than two cells. Such scenario is better recapitulated by a model with increased cooperativity (c=5.5) (Fig. 4C). For these three datasets, the frequency of single cells was not reported due to technical limitations. Our model predicts that single cell migration is the prevalent migration mode in these three cases, with a fraction of 92%, 84% and 55% of single CTCs, respectively (supplementary table 1). A mouse model of breast cancer [22] shows a broad distribution with large clusters of 6-8 cells (Fig. 4D). While our model can qualitatively capture the transition toward a more clustered migration, the probability to observe clusters remains relatively constant within a certain range of size in the experimental dataset (Fig. 4D). A further increase of cell cooperativity, however, gives rise to a different regime where small clusters become more probable than single cells (c=8.5, Fig. 4E). The onset of this single cell-cluster transition recapitulates well the size distribution measured in a mouse model of myeloma [23] that peaks for a cluster size of n=4 cells (Fig. 4E). In this case, single cells and small clusters of n=2 cells could not be measured experimentally, but our model predicts those to account approximately for only 3% and 8% of the total CTC fraction, while larger clusters of 3-5 cells account for about 68% of the total CTC fraction (supplementary table 1). CTC clusters detected in human glioblastoma [24] (Fig. 4F) are mapped onto an even higher cooperativity (c=9). Similar to the case of breast cancer (Fig. 4D), our model qualitatively captures the shape of the distribution but underestimates the frequency of large clusters composed by 7-8 or more cells (Fig. 4F). Finally, another mouse model of breast cancer [3] is characterized by a typical cluster size of 5-10 cells. This size distribution can be mapped onto a very high cooperativity index (c=10) without increasing the rate of EMT (Fig. 4G).

**Figure 4.**
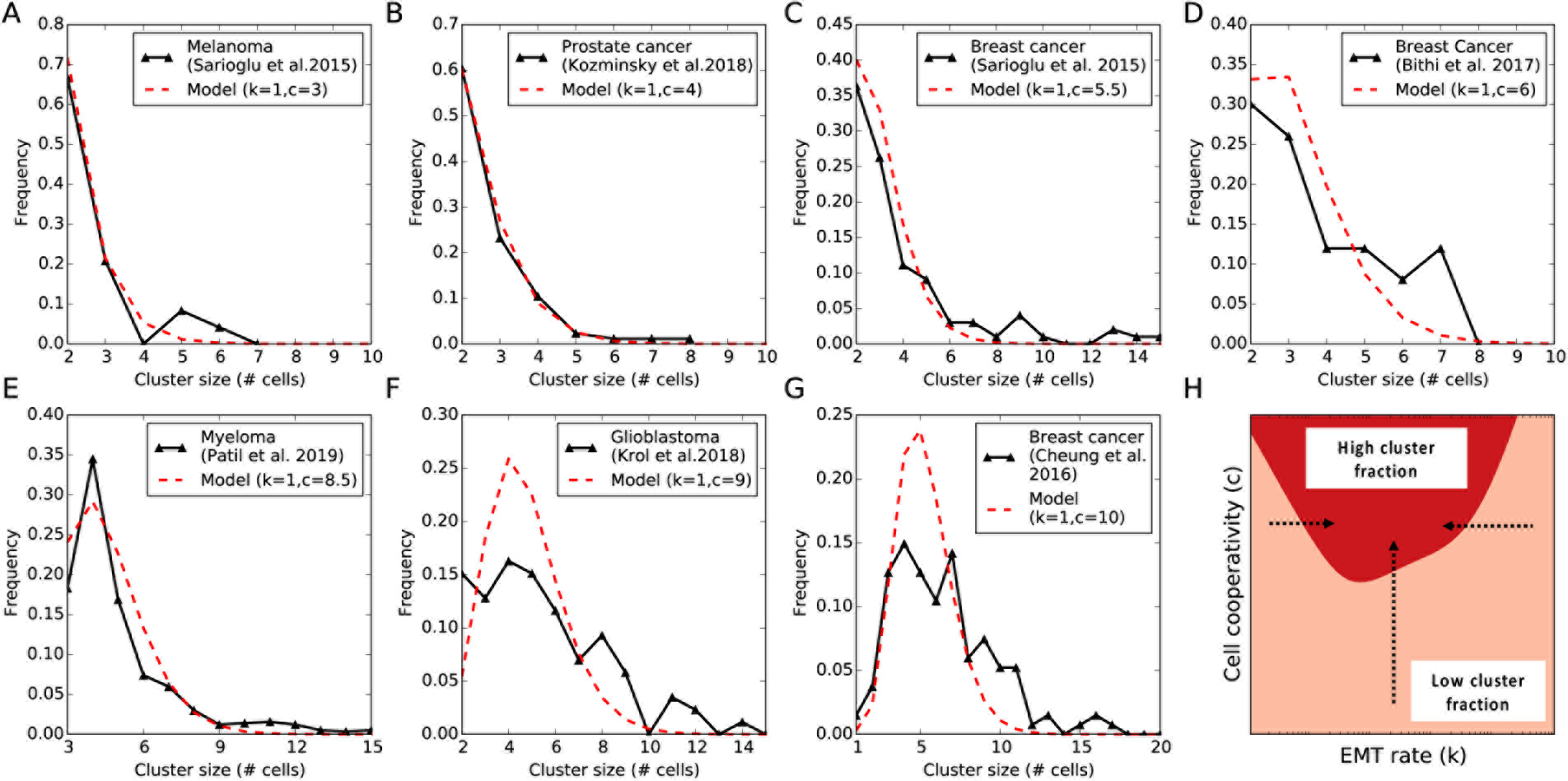
The single cell-cluster migration transition recapitulates cluster size distributions across several cancer types. CTC cluster size distributions from various experiments and model’s fit for a fixed EMT rate (k=1) and increasing cooperativity index (c) **(A)** Human melanoma(adapted from Sarioglu et al. [21]. **(B)** Human prostate cancer adapted from Kozminsky et al. [15]. **(C)** Human breast cancer adapted from Sarioglu et al. [21]. **(D)** A mouse model of breast cancer adapted from Bithi et al. [22]. **(E)** A mouse model of myeloma adapted from Patil et al. [23]. **(F)** Human glioblastoma adapted from Krol et al. [24]. **(G)** A mouse model of breast cancer adapted from Cheung et al. [3]. For datasets A-B-C-E-F, single cells were not measured; for dataset D, single cells and clusters of size n=2 cells were not measured. The original datasets were normalized to help comparison with the model. **(H)** Qualitative representation of the transition pathways from single cell migration to cluster cell migration in the (k, c) parameter space. This diagram depicts a ‘binarized’ version of Fig. 2C where the parameter space is differently colored depending on whether the cluster escape fraction is smaller (orange) or larger (red) than 0.5.

These datasets were reproduced with a fixed EMT rate and increasing cooperativity. However, this is not the only combination in the two-dimensional parameter space of (k, c) that can reproduce these distributions. Indeed, several combinations of (k, c) can quantitatively reproduce these distributions (Fig. S5). In summary, a progressing tumor can access the cluster-based dissemination mode by (i) increasing the migration cooperativity at fixed EMT rate, (ii) increasing/decreasing its EMT rate from an extreme value and large enough cooperativity, or (iii) through a combination of these two pathways. (Fig. 4H).

### A model of only partial, and not complete EMT, better recapitulates collective migration

The migration model with 3 cell states (E, E/M, M) could not quantitatively capture some features of the CTC cluster distributions in the datasets where migration is dominated by clusters, including (i) broad distributions where CTC clusters of different sizes have similar frequencies (see Fig. 4D, 4F) and (ii) the frequency of large CTC clusters of more than 8-10 cells (see Fig. 4F-G). Different molecular mechanisms, however, can interfere with the multiple steps of EMT, stabilize hybrid E/M phenotype(s) and give rise to more CTC clusters [25–28]. To address this scenario, we separately consider a rate of partial EMT (E→E/M transition) and a rate of complete EMT (E/M→M transition) (Supplementary Information H).

A fast partial EMT, coupled with a slow complete EMT, generates a high fraction of hybrid E/M cells in the lattice (Fig. 5A). A slow partial EMT, however, results in an epithelial lattice, while the tissue is mostly mesenchymal when both the processes are fast (Fig. S6A-B). Therefore, an increased rate of complete EMT (*k*_*Complete EMT*_) does not favor a large fraction of cells disseminating as clusters (Fig. S6C). Indeed, a high expression of ‘phenotypic stability factors’ that can stabilize hybrid epithelial/mesenchymal phenotype(s) and increase the mean residence times in these phenotypes by preventing them from undergoing a complete EMT (i.e. reduce *k*_*Complete EMT*_ effectively) have been correlated with higher aggressiveness and worse patient survival in multiple cancer datasets [26–29], corroborating the evidence of clusters being the primary ‘bad actors’ of metastasis.

**Figure 5.**
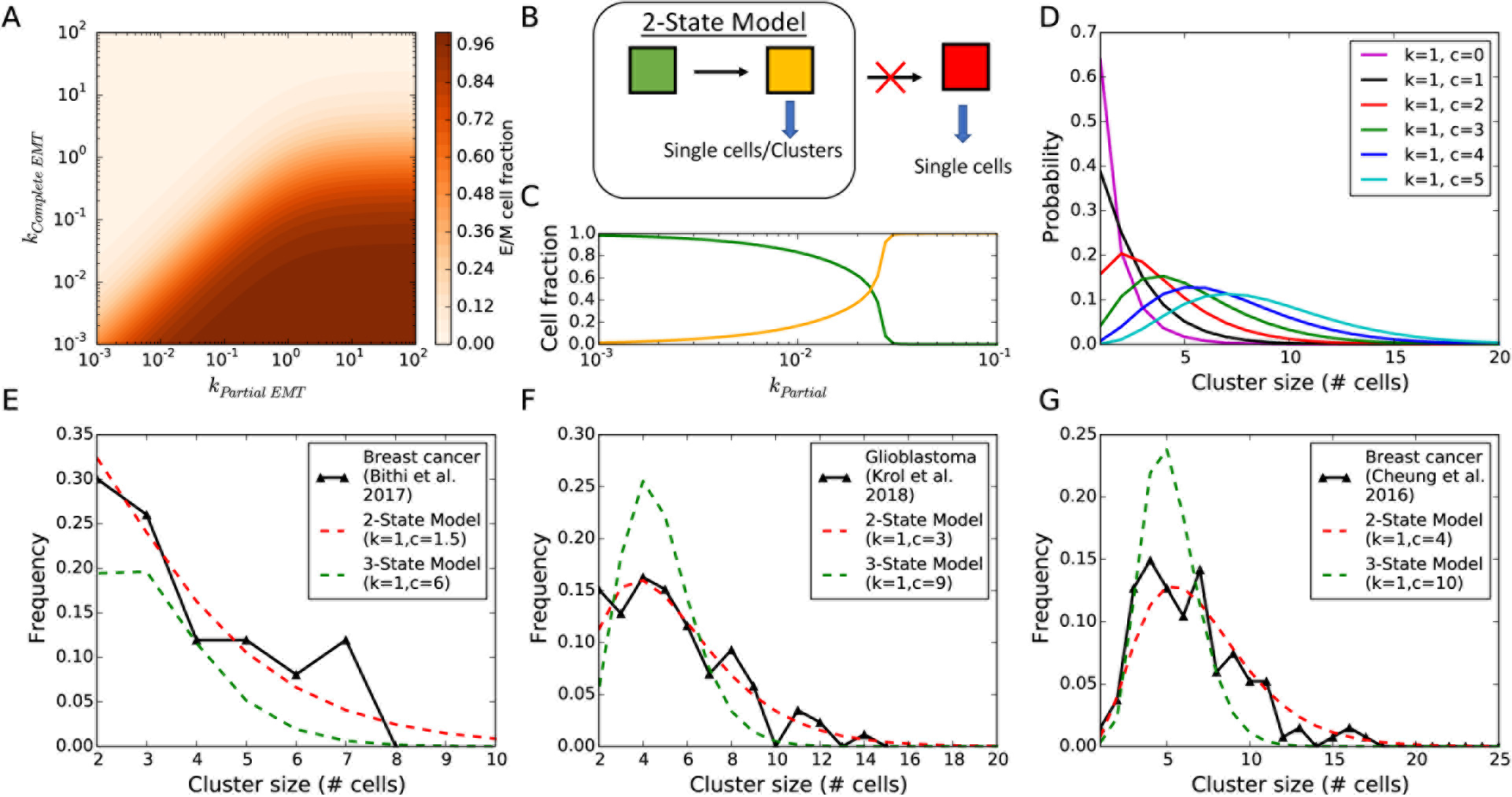
Partial EMT reproduces collective migration. **(A)** Fraction of hybrid E/M cells as a function of the rates of partial EMT (*k*_*Partial EMT*_) and complete EMT (*k*_*Complete EMT*_) for c=1. **(B)** Limit case where the rate of complete EMT is very small. **(C)** Fractions of E cells (green) and E/M cells (yellow) in the 2-State model as a function of *k*_*Partial EMT*_. **(D)** Cluster size distribution in the 2-state model for increasing values of migration cooperativity (c) at fixed *k*_*Partial EMT*_(denoted as ‘k’ for simplicity). **(E-F-G)** Experimental cluster size distributions with large clusters and best fit as predicted by the 2-state model (red dashed) and by the 3-state model (green dashed, same as Fig. 4).

In the limit where the rate of progression to a complete EMT state is very slow (*k*_*Complete EMT*_ ≪ *k*_*Partial EMT*_), the model is restricted to the epithelial state and the hybrid E/M state (Fig. 5B). In this 2-state limit, a large rate of partial EMT ensures that almost all cells are hybrid E/M (Fig. 5C). In this particular situation, cells can still migrate more solitarily or collectively depending on the cooperativity index (c). Without cooperativity, only single cells and small clusters are observed, while a higher cooperativity results in larger clusters (Fig. 5D).

Strikingly, this 2-state model correctly captures the broad size distributions observed in breast cancer and glioblastoma [22,24] (Fig. 5E-F) and the collective invasion with large clusters >10 cells reported in breast cancer [3] (Fig. 5G). This picture is consistent with the fact that, at least in breast cancer, a conserved epithelial program, rather than a complete EMT, seems to be required for collective invasion [9]. Additionally, we note that a much smaller (and possibly more meaningful from a biological standpoint) value of cooperativity (c) is required to reproduce large clusters in the 2-state model (see definition of c in Supplementary Information A).

### Multiple intermediate hybrid states facilitate cluster dissemination and heterogeneity

Recent experiments suggest that multiple hybrid states with mixed epithelial and mesenchymal traits can exist with distinct epigenetic landscapes and morphological features [7,30,31].

Therefore, we generalize the model to an arbitrary number (n) of hybrid states. It is assumed that an epithelial cell transitions through the n intermediates in an ordered fashion to become mesenchymal. Furthermore, the intermediate states are orderly ranked, and every transition (*E*/*M*_*i*_ → *E*/*M*_*i*+1_) makes the cell less epithelial and more mesenchymal, i.e. cell becomes less and less adhesive. In the model, cells in different intermediate states can still migrate together as clusters (Supplementary Information - I).

Increasing the number of intermediate states increases the cluster escape fraction and the probability to observe rare events of large clusters (Fig. 6A-B). Moreover, multiple intermediate states enable cluster-based dissemination with a higher fraction of CTC clusters at a much lower cooperativity index, compared with the case of only one intermediate state (Fig. S7A).

**Figure 6.**
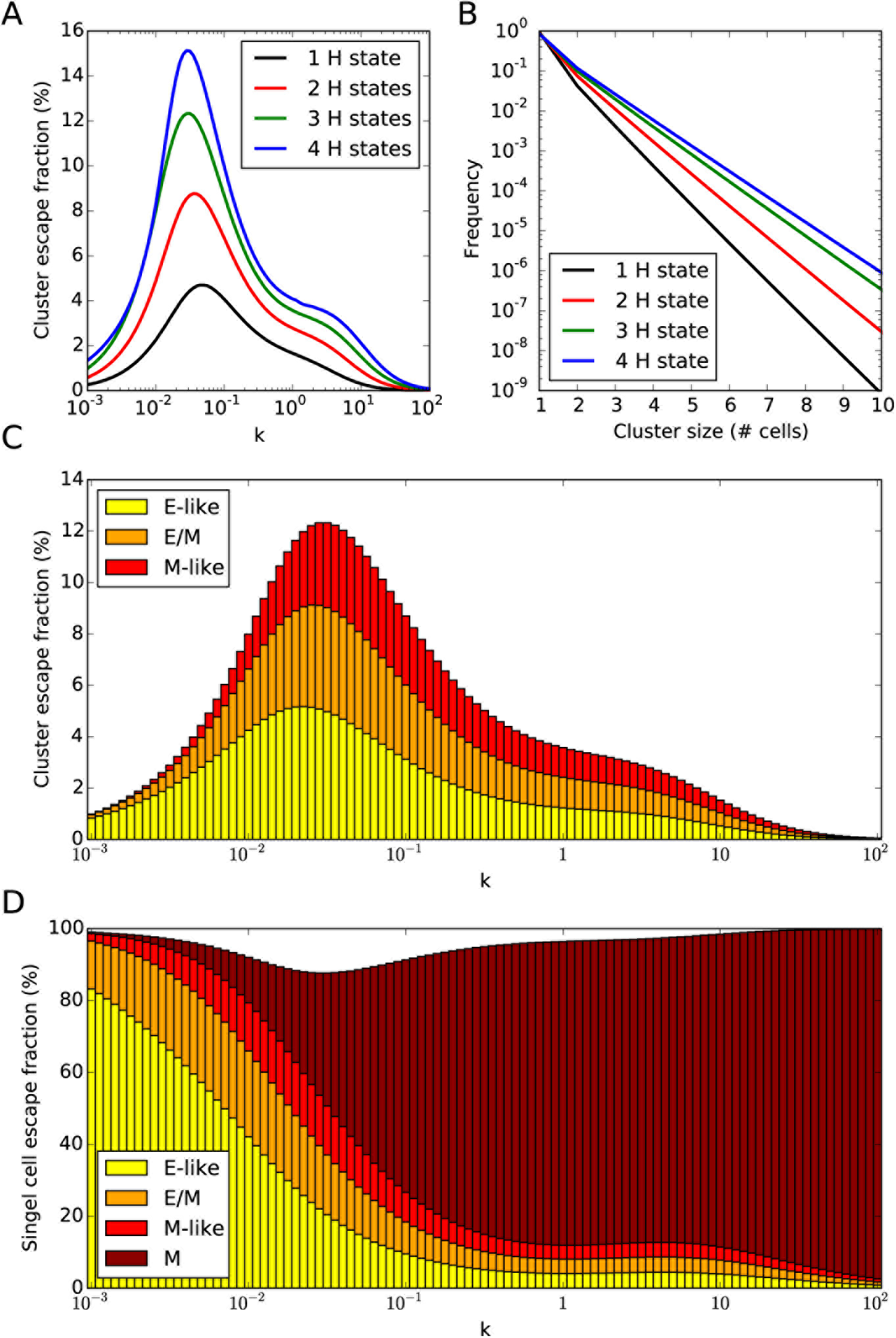
Multiple intermediate hybrid E/M states facilitate cluster dissemination and heterogeneity. **(A)** Cluster escape fraction as a function of the EMT rate (k) and **(B)** cluster size distribution for models with an increasing number of intermediate hybrid E/M states. **(C)** Cluster escape fraction as a function of k in the model with 3 intermediate states. Color bars show the average cluster composition in terms of fractions of E-like, E/M and M-like cells. **(D)** Single cell escape fraction as a function of k in the model with 3 intermediate states. Color bars show the fraction of single E-like, E/M, M-like and Mesenchymal migrating cells (c=1 for all panels).

Moreover, multiple intermediate states result in a more heterogeneous cluster composition. In the case of three intermediate states, called E-like, E/M and M-like here, clusters are a mixture of cells in the three intermediate states. (Fig. 6C). This heterogeneity has been well-reported in both clusters cultivated *in vitro* and circulating tumor clusters *in vivo*, where staining revealed that (i) only certain cells in the cluster express epithelial or mesenchymal markers and (ii) certain cells robustly express both epithelial and mesenchymal markers [2,32–34].

Similarly, multiple intermediate states give rise to a spectrum of single migrating cells with different EMT traits (Fig. 6D). If EMT is slower than the cell migration (*k* ≪ 1), most cells migrate as soon as they transition to an E-like state. Therefore, in this case, most of the migrating cells lean toward the epithelial end of the EMT spectrum (Fig. 6D, left region). Conversely, for a faster EMT (*k* ~ 1), most migrating cells lean toward the mesenchymal end of the EMT spectrum (Fig. 6D, right region). The heterogeneous composition of cluster and single cell states is conserved even for variation of the migration cooperativity parameter (c) (Fig. S7B-C). This heterogeneity has been well reported experimentally and could be informative of the tumor progression and response to treatments. We compared the model’s predictions with the CTC cell fractions measured in the bloodstream of breast cancer patients [35] (Supplementary Information - J). Notably, patients that responded positively to treatment maps onto smaller EMT rate in the model after treatment, while patients that underwent tumor progression maps onto a larger EMT rate after treatment (Fig. S8).

Multiple intermediate states generate more clusters similar to the slower transition of a single intermediate state (see previous section). However, different cell states can potentially cooperate and perform different functions, hence increasing the survival probability of a heterogeneous CTC cluster.

A different multistate EMT scenario would consider parallel pathways from E to M that pass through different intermediate states, rather than a ‘linear chain’ of states (Fig. S9A). These two different scenarios present slight differences in terms of cell state fractions as well as CTC cluster fractions (Fig. S9B-C). To discern between these different scenarios in real tumors, however, more data will be necessary to clarify whether (i) the EMT spectrum is truly a set of discrete state or a more continuum range and (ii) whether different hybrid states serve different purposes, such as EMT and its reverse, MET.

## Discussion

We propose a model that couples cell migration with the phenotypic plasticity driven by the epithelial-mesenchymal phenotypic transition. The model predicts a transition from a migratory regime mostly dominated by single mesenchymal cells to a regime mostly dominated by hybrid Epithelial/Mesenchymal cell clusters. This transition depends on the rate at which cells undergo EMT and on the level of cooperativity that quantifies the adhesion between collectively migrating cells and the interaction between cancer cells and the extra-cellular matrix (ECM). This simple biophysical model can quantitatively reproduce different cluster size distributions observed in the bloodstream of patients and/or in experimental mouse models. These experimental datasets exhibit a tremendous variability, ranging from cases where large clusters are highly improbable to cases of collective migration [3,15,21–24], all of which can be captured by the model for varying parameter combinations.

Nonetheless, we acknowledge multiple limitations to our model. First, cancer cells can proliferate, hence introducing an additional modulation to cell patterning. Second, cell intravasation and death in circulation may depend on the cluster size [36]. Third, additional mechanisms of aggregation can modulate the size of these clusters, such as collisions in the bloodstream [37]. Fourth, CTC clusters may contain non-cancerous cells such as neutrophils [38]. Finally, the abovementioned datasets have been acquired in different systems and with different experimental techniques, which might have affected the accuracy in detection of CTC clusters. With these caveats being raised, these results support the idea that multiple aspects of cancer cell migration are highly conserved across cancer types, and microenvironmental, case-specific factors might set the conditions for a more individual or collective migration by regulating cancer cell heterogeneity in terms of EMT and migration. Supporting this picture, it has been recently shown that multiple breast cancer cell lines with varying origin and morphological features can switch from solitary to collective migration by manipulating the physical properties of the extracellular medium *in vitro* [12,13].

The existence of one or more hybrid E/M phenotypes capable to invade while maintaining cell-cell adhesion has been proposed as a possible explanation for the different modes and aggressiveness of cancer invasion [39]. Supporting this hypothesis, hybrid E/M phenotype(s) have been associated with stemness and tumor-initiation ability [40–44], while inducing a complete EMT has been shown to even restrict tumor aggressiveness and metastatic potential in some cases [45–48]. While our model of complete EMT could correctly recapitulate the size distribution of CTC clusters in contexts where the migration is mostly solitary, a model limited to only partial EMT could quantitatively reproduce cases of both solitary and collective migration. Therefore, although a complete EMT can take place in certain types of cancer, a partial EMT might be sufficient to explain the different modes of invasion, and seems necessary for collective migration.

Moreover, we show that multiple, instead of a single, hybrid E/M phenotypes give rise to heterogeneous clusters composed by cells with different positions along the ‘EMT spectrum’. This heterogeneity might enable co-operation among different phenotypes in executing various aspects of metastasis, hence increasing the survival probability of clusters in circulation. For instance, completely mesenchymal CTCs are associated with a more favorable survival outcome in breast cancer [49], and completely mesenchymal cells lose their metastatic potential in squamous cell carcinoma [50].

This framework offers a platform that could be integrated in the future with more explicit models of intracellular regulatory dynamics and cell-cell and cell-environment mechanical interactions *en route* to a better understanding of cancer metastasis.

The authors declare no potential conflict of interest

## Acknowledgements

The authors thank Prof. Herbert Levine and Prof. Andrew Ewald for their valuable suggestions.

