## Supplementary information and figures for "A biophysical model uncovers the size distribution of migrating cell clusters across cancer types"

### Details on Mathematical Model

**A - Model of the escape rate**
M cells escape from the lattice only as single cells with a basal escape rate $k_{ESC}$. Conversely, H cells can escape either as single cells or as multi-cellular clusters. The breaking of bonds between cells are considered as independent stochastic processes defined by the probabilities to break an E-H bond and an H-H bond ($p_E$ and $p_H$, respectively). Since the current work will only be concerned with the average properties of the system, we introduce these terms as modulation to the cluster escape rate rather than by simulating stochastic events. In other words, the escape rate of a subcluster composed by m H-cells enclosed within a larger cluster composed by s H-cells is:

$$K(m,s) = m^c {p_E}^{n_E} {p_H}^{n_H} \qquad (1)$$

In this formula, c is a migration cooperativity index, and $n_E$ and $n_H$ are the average number of nearest neighbor cells of the cluster in an E state and an H state, respectively, and are calculated exactly in the section E below. In general, this rate depends on the dimension and connectivity of the considered lattice (see Supplementary figure 1A for an example on a 2D square lattice). In the limit case of a single mesenchymal (M) cell, m=1 and there are no adhesion bonds, therefore eq. (1) naturally relaxes to the basal escape rate $k_{ESC}$.

**B - Flux of escaping clusters**
In a generic lattice with arbitrary dimension and connectivity, the flux of escaping clusters of H cells per unit time is quantified by considering every cluster in the lattice, every possible subcluster that could arise from any of those clusters, and the escape rate of any subcluster given by eq. 1. This approach, however, becomes computationally too expensive when considering large lattices. Therefore, we consider a mean-field approximation for the cluster flux:

$$\Theta(\rho_H) = \sum_{s=1}^{+\infty} p_s(\rho_H) \sum_{m=1}^{s} n(m,s)\, K(m,s) \qquad (2)$$

In this expression, the first summation considers the probability to find clusters of a certain size s ($p_s(\rho_H)$) for a certain fraction of H cells in the lattice ($\rho_H$). The second summation considers the relative frequencies ($n(m,s)$) and escape rates ($K(m,s)$) of subclusters of size m<s that could arise from a given cluster of size s. The analytical expression of the cluster size distribution $p_s(\rho_H)$, however, is not known in any dimension $D \geq 2$. Therefore, we consider cells arranged in an infinite one-dimensional chain. In this case, $p_s(\rho_H) = {\rho_H}^s (1-\rho_H)^2$ from 1D percolation theory (13), and the relative frequencies of subclusters ($n(m,s)$) and number of nearest neighbors ($n_E$, $n_H$) can be computed analytically as well (see sections below).

**C - Model setting – an 'effective-2D' approximation**
We consider the cells to be arranged in an infinite one-dimensional chain. This choice allows to calculate analytically the distribution of clusters as a function of the fraction of H cells in the chain ($\rho_H$) (13). To make the model more realistic from a biological standpoint (i.e. tumor-stroma interfaces are typically 2D surfaces), we consider an "effective-2D" approximation. In the effective-2D setting, the infinite 1D chain is in contact with two replicas that have the same fractions of E, H and M cells (Fig. S1C). This assumption reflects the fact that, in a 2D or 3D tissue, larger clusters have more nearest neighbors, and so more bonds to break to migrate. Moreover, an explicit simulation on a 2D lattice would rapidly become computationally expansive when considering lattices with a number of cells N at least one order of magnitude larger than the typical cluster sizes that will be considered (N≈100). In fact, calculating the escape flux of clusters as defined in section B would require enumerating at any given time all the clusters of H cells and all the subclusters that can arise from breaking of these clusters. Compared to a 2D lattice model, the effective-2D approximation underestimates the migration of large clusters for two main reasons. First, for the same fraction of H cells in the chain ($\rho_H$), the probability to form large clusters is larger in a 2D lattice (supplementary figure 1B). Second, clusters in the effective-2D model are linear chains. Therefore, their

number of nearest neighbors, and so the number of bonds to break, is maximized as compared to a 2D lattice (supplementary figure 1C).

**D - Frequency of subclusters in a one-dimensional chain**

In a one-dimensional chain, a cluster of s cells can only be a linear chain. The number of possible subclusters of size m (m<s) that can be formed from the cluster of size s is:

$$N(m,s) = s - m + 1 \qquad (3)$$

The relative frequency of subclusters of a given size m equals the number of subclusters of size m divided by the total number of subclusters of any size that can be formed from the original cluster:

$$n(m,s) = \frac{N(m,s)}{\sum_{m=1}^{s} N(m,s)} = \frac{s-m+1}{\frac{s(s+1)}{2}} \qquad (4)$$

**E - Number of nearest neighbors in mean field approximation**

$n_E$, $n_H$ and $n_M$ are the number of nearest neighbors of the cluster that are in an E, H, and M state, respectively. In the one-dimensional chain, each cluster or subcluster has always 2 nearest neighbors, independently from its size. Therefore, on average, $n_E = 2\rho_E/(\rho_E + \rho_M)$ nearest neighbors and $n_M = 2\rho_M/(\rho_E + \rho_M)$. A subcluster of size m enclosed within a cluster of size s, however, can have nearest neighbors in the H state (see supplementary Fig. 1A for a visual representation). Therefore, for the subcluster of size m, two scenarios are possible. Out of the $s - m + 1$ possible subclusters, $s - m - 1$ are completely enclosed within the main cluster, and their nearest neighbors are therefore 2 H cells. The remaining 2 subclusters of size m share 1 bond with an H cell within the cluster and another bond with either an E cell or an M cell with probabilities $\rho_E/(\rho_E + \rho_M)$, $\rho_M/(\rho_E + \rho_M)$, respectively.

In the effective-2D variation of the model, a cluster of size s has 2 nearest neighbors from the main lattice and s additional neighbors from each one of the two replicas, for a total of 2+2s nearest neighbors. In this case, a cluster can have H nearest neighbors as well. Similarly, a subcluster of size m has 2+2m nearest neighbors. The additional neighbors resulting from the effective-2D correction are in an E, H or M state with probabilities $\rho_E$, $\rho_H$, $\rho_M$, respectively. In any dimension equal or larger than 2, the cluster size distribution is not known, and a simulation is required.

**F - Temporal dynamics and steady state of the model**

The temporal dynamics of E, H, and M cell fractions ($\rho_E$, $\rho_H$, $\rho_M$) in the infinite, effective-2D lattice are described by ordinary differential equations that consider EMT events, escape of H and M cells, and replacement by new E cells:

$$\frac{d\rho_E}{dt} = -k_{EMT}\,\rho_E + k_{ESC}\Theta(\rho_H) + k_{ESC}\rho_M \qquad (5a)$$

$$\frac{d\rho_H}{dt} = +k_{EMT}\,\rho_E - k_{EMT}\rho_H - k_{ESC}\Theta(\rho_H) \quad (5b)$$

$$\frac{d\rho_M}{dt} = +k_{EMT}\rho_H - k_{ESC}\rho_M \qquad (5c)$$

To model EMT transitions, the epithelial cell population ($\rho_E$) has an outward flux $-k_{EMT}\,\rho_E$ due to partial EMT transition; the H cell population ($\rho_H$) increases with rate $k_{EMT}\,\rho_E$ due to partial EMT but decreases with rate $-k_{EMT}\rho_H$ due to transition to a complete EMT; finally, the mesenchymal cell population ($\rho_M$) increases with rate $k_{EMT}\rho_H$. Here, $k_{EMT}$ is a basal rate of EMT with units of inverse time. Moreover, to model cell migration, the H cell population decreases with rate $-k_{ESC}\Theta(\rho_H)$, where $k_{ESC}$ is a basal rate constants for cell escape with units of inverse time and $\Theta(\rho_H)$ is the instantaneous dimensionless flux of escaping clusters defined in section (B). Since mesenchymal cells escape only as single cells with rate $k_{ESC}$, the loss term for mesenchymal cell population is simply $-k_{ESC}\rho_M$. The escape rate terms $k_{ESC}\Theta(\rho_H)$, $k_{ESC}\rho_M$ increase the epithelial cell population to conserve the total cell density and model replacement by new epithelial cells upon migration.

Since the current work will only be concerned with steady state properties, we define a dimensionless time $t' = k_{ESC}t$. Therefore, the model is described by a single dimensionless rate $k = k_{EMT}/k_{ESC}$ that represents the ratio between the basal rates of EMT and escape:

$$\frac{d\rho_E}{dt} = -k\,\rho_E + \Theta(\rho_H) + \rho_M \qquad (6a)$$
$$\frac{d\rho_H}{dt} = +k\,\rho_E - k\,\rho_H - \Theta(\rho_H) \quad (6b)$$
$$\frac{d\rho_M}{dt} = +k\,\rho_H - \rho_M \qquad (6c)$$

In the main text, we will refer to k simply as the rate of EMT since it appears in the EMT reaction terms, while the basal escape rate will always be set to unit value. Setting Eqs. 6 to zeros provide the steady state cell fractions ($\rho_E{}^{(ss)}$, $\rho_H{}^{(ss)}$, $\rho_M{}^{(ss)}$), from which the steady state distribution of escaping clusters can be reconstructed (see section below).

**G – Steady state cluster size distribution**

The probability distribution of escaped clusters as a function of cluster size, or cluster size distribution, can be reconstructed as follows. In the dimensionless model, the total escape flux of clusters of fixed size s is:

$$F(s) = \sum_{s'>s} p_{s'}(\rho_H) \sum_{m=1}^{s} n(m,s')\,k(m,s')\delta_{ms} = \sum_{s'>s} p_{s'}(\rho_H) n(s,s')\,k(s,s') \qquad (7)$$

Where the condition $s' > s$ considers that a subcluster of size s can arise from clusters of size s' larger or equal than s, and the Kronecker's delta $\delta_{ms}$ ensures that only subclusters of size s are considered. Finally, to compute the cluster size distribution $f(s)$, one must consider that the escape flux of single cells (s=1) receives the additional contribution of M cells that can only migrate as single cells:

$$f(s) = \begin{cases} \dfrac{\rho_M + F(1)}{\rho_M + \Theta(\rho_H)}, & s = 1 \\ \dfrac{F(s)}{\rho_M + \Theta(\rho_H)}, & s > 1 \end{cases} \qquad (8)$$

**H - Variation of EMT rates and 2-state model**

The rates pf partial EMT and complete EMT are independent parameters, so that:

$$E \xrightarrow{k_{Partial\,EMT}} H \xrightarrow{k_{Complete\,EMT}} M$$

In the limit where $k_{Complete\,EMT}{\sim}0$, the model has only the E and H states, with the following reactions:

$$E \xrightarrow{k_{Partial\,EMT}} H$$
$$H \xrightarrow{Escape} E$$

**I - Multiple intermediate states**

For a generic case of n IS, it is assumed that an E cell must transition through all ISs in an ordered fashion to become mesenchymal (i.e. a 'linear chain model'):

$$E \xrightarrow{k_{EMT}} H_1 \xrightarrow{k_{EMT}} H_2 \xrightarrow{k_{EMT}} \dots \xrightarrow{k_{EMT}} H_{n-1} \xrightarrow{k_{EMT}} H_n \xrightarrow{k_{EMT}} M$$

Cells in different IS can form clusters and escape together. Therefore, in a model with n IS, each IS *i* is characterized by a set of n probabilities: the probability of breaking a bond between an epithelial cell and a cell in the $i^{th}$ IS, plus all the probabilities of breaking bonds between a cell in the $i^{th}$ IS and a cell in any other IS. To estimate these probabilities, it is assumed that the cell states are ordered on an adhesion scale A, so that the E state has a maximal adhesion factor $A_E$=1 and the M state has a minimal adhesion factor $A_M$=0. All IS are placed in an ordered and linear fashion in (0,1), so that $A_{i-1}>A_i>A_{i+1}$. With this scale, a pairwise adhesion factor $A_iA_j$ can be defined for two cells in IS *i* and *j*, respectively. This factor is bound in [0,1], and can be interpreted as the probability of maintaining the bond. The probability of breaking the bond can be thus defined as the complementary probability $p_{ij}$=1- $A_iA_j$. When applied to the simple case of one IS, this generalization replicates the values of $p_E$ and $p_H$ with $A_E$=1, $A_H$=0.5, $A_M$=0.

To model the case of multiple EMT pathways passing through different intermediate states of partial EMT, we consider two intermediates ($H_1$ and $H_2$) such that EMT can take place through the $E \rightarrow {H_1}^p \rightarrow M$ or the $E \rightarrow {H_2}^p \rightarrow M$ parallel pathways. We assume a similar rate k for all the transition ns. Similar to the case of ordered EMT through multiple intermediates, neighbors in the ${H_1}^p$ and ${H_2}^p$ state can form clusters together and migrate, and ${H_1}^p$ and ${H_2}^p$ differs only for their different adhesion with neighbors, modeled with different probability to break bonds. The cell state ${H_1}^p$ is characterized by the probabilities to break bonds with E, ${H_1}^p$ and ${H_2}^p$ ($p_{E1}$, $p_{H1}$ and $p_{H12}$, respectively), while the cell state ${H_2}^p$ is characterized by ($p_{E2}$, $p_{H2}$ and $p_{H21} = p_{H12}$). Since this model of 'parallel pathways' will be compared to the 'linear chain model' with two ISs, we assume similar probabilities to break bonds for the intermediates ${H_1}^p$ and ${H_2}^p$ than those of the ISs $H_1$ and $H_2$ of the 'linear chain model' (namely: $p_{E1} = 1/3$, $p_{H1} = 5/9$ and $p_{H12} = 7/9$; $p_{E2} = 2/3$, $p_{H2} = 8/9$ and $p_{H21} = p_{H12} = 7/9$).

**J – K-score of patient datasets**

In the considered datasets, CTCs were classified as E, E>M, E=M, M>E, M. These fractions were compared to the fractions of escaping single cells predicted by the model. Since the E state is not motile in the model, we considered a case with 4 intermediate hybrid states. The fractions of single migrating cells in the 4 intermediates and in the mesenchymal state were compared to the 5 CTC fractions from the dataset, hence defining a distance between the dataset and the model. The k score is the value of EMT rate (k) that minimizes the dataset between the experimental dataset and the model's prediction at a fixed cooperativity c=1.

### Supplementary Figures and Tables

**Supplementary figure 1.** The effective-2D model underestimates the escape of large clusters as compared to a 2D model.

**Supplementary figure 2.** Parameter sensitivity of the effective-2D model.

**Supplementary figure 3.** CTC cluster detection ranges from independent experiments are mapped onto different regions of the parameter space.

**Supplementary figure 4.** Comparison between 1D, effective-2D and square-2D models.

**Supplementary figure 5.** Experimental cluster size distributions are mapped onto different regions of the parameter diagram.

**Supplementary figure 6.** Control of partial and complete EMT modulates the fraction of E, H, and M cells in the lattice.

**Supplementary figure 7.** The cooperative migration conserves the heterogeneous composition of clusters and single cells.

**Supplementary figure 8.** CTC percentages of breast cancer patients map onto different EMT rates of the model.

**Supplementary figure 9.** Comparison between different multistate EMT models.

**Supplementary Table 1.** Cluster escape flux percentage as a function of cluster size.

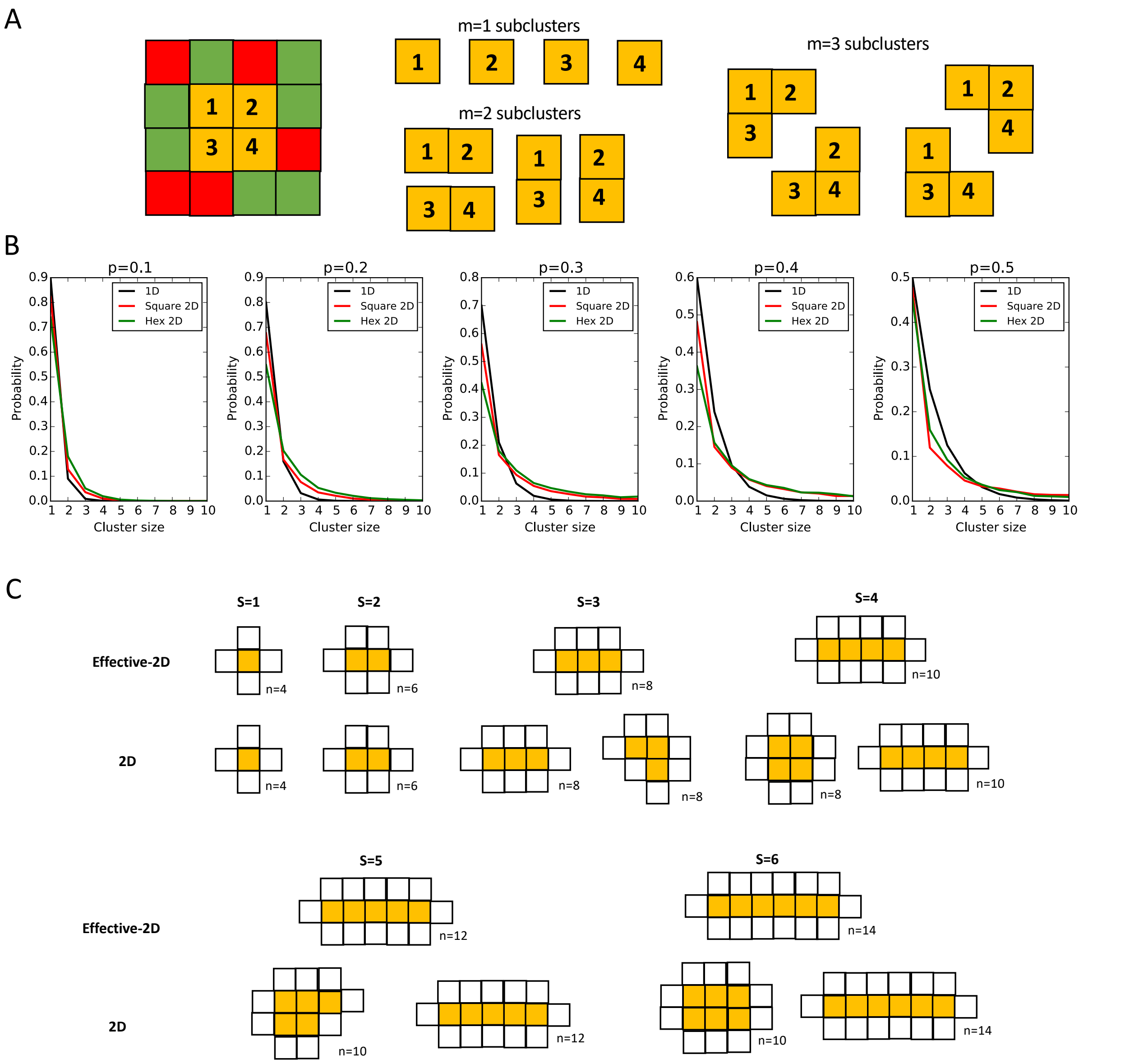


Supplementary figure 1. **The effective-2D model underestimates the escape of large clusters as compared to a 2D model. (A)** In this example on a 2D square lattice, a cluster of s=4 H cells (marked as yellow) is present {1,2,3,4}. Additionally, there are 4 subclusters of m=1 cells, 4 subclusters of m=2 cells ({1,2}, {1,3}, {2,4}, {3,4}), and 4 subclusters of m=3 cells ({1,2,3}, {1,2,4}, {2,3,4}, {1,3,4}). In total, there are 1+4+4+4=13 possible escaping structures. Each of these subclusters can escape with a rate K(m,s=4) given by eq. 1. The dependence on the overall cluster size (s=4) arises because the adhesion term in eq. 1 depends on the cell state of the nearest neighbors of the cluster. **(B)** Distribution of clusters as a function of size for varying fractions of H cells. Note that this is the distribution of clusters that are formed in the lattice (i.e. the conventional cluster size distribution considered in lattice models), and not the distribution of escaped clusters. Each panel compares the distribution for 1D chain, square-2D and hexagonal-2D lattices. Large clusters are more probable in the 2D distributions. **(C)** The effective-2D model correctly reproduces the number of nearest neighbors (n) of small clusters ($s \leq 3$) as compared to a 2D square lattice, but large clusters ($s \geq 4$) can be arranged in more and more compact configurations in a square 2D lattice as compared to the effective-2D model.

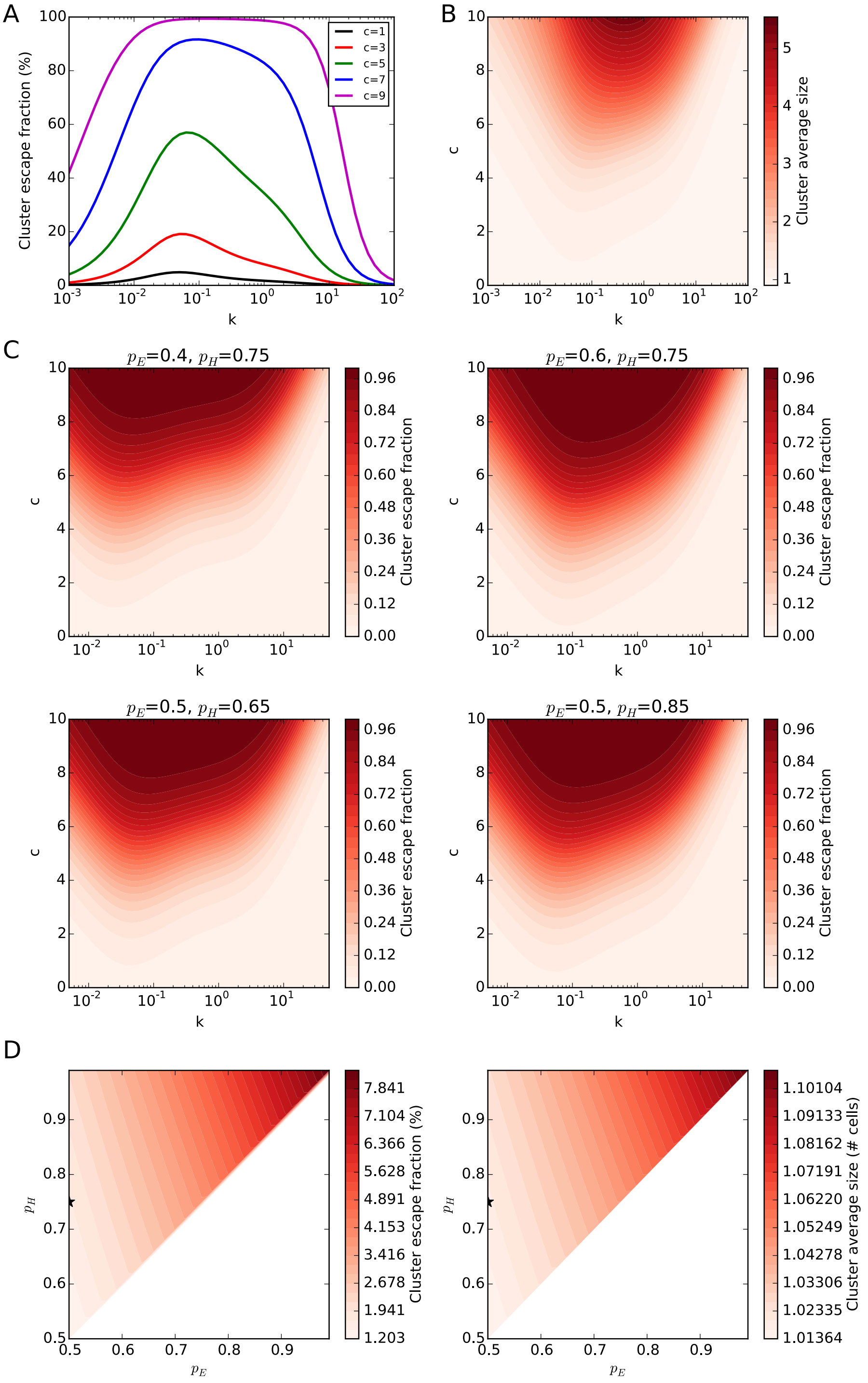


Supplementary figure 2. **Parameter sensitivity of the effective-2D model. (A)** Cluster escape fraction as a function of the EMT rate (k) for different levels of cooperativity (c). **(B)** Cluster average size as a function of the EMT rate (k) and the cooperativity (c). The average considers also single cells. **(C)** Cluster escape fraction as a function of k and c for different combinations of the probabilities of breaking an E-H bond ($p_E$) and an H-H bond ($p_H$). The reference values for Figure 2C are pE=0.5, pH=0.75. **(D)** Cluster escape fraction and cluster average size as a function of $p_E$ and $p_H$ (k=1, c=1). $P_H$ is always assumed to be larger than $p_E$, therefore only the half of the parameter space where $p_H$>$p_E$ is studied. The start indicates control values used in the main text.

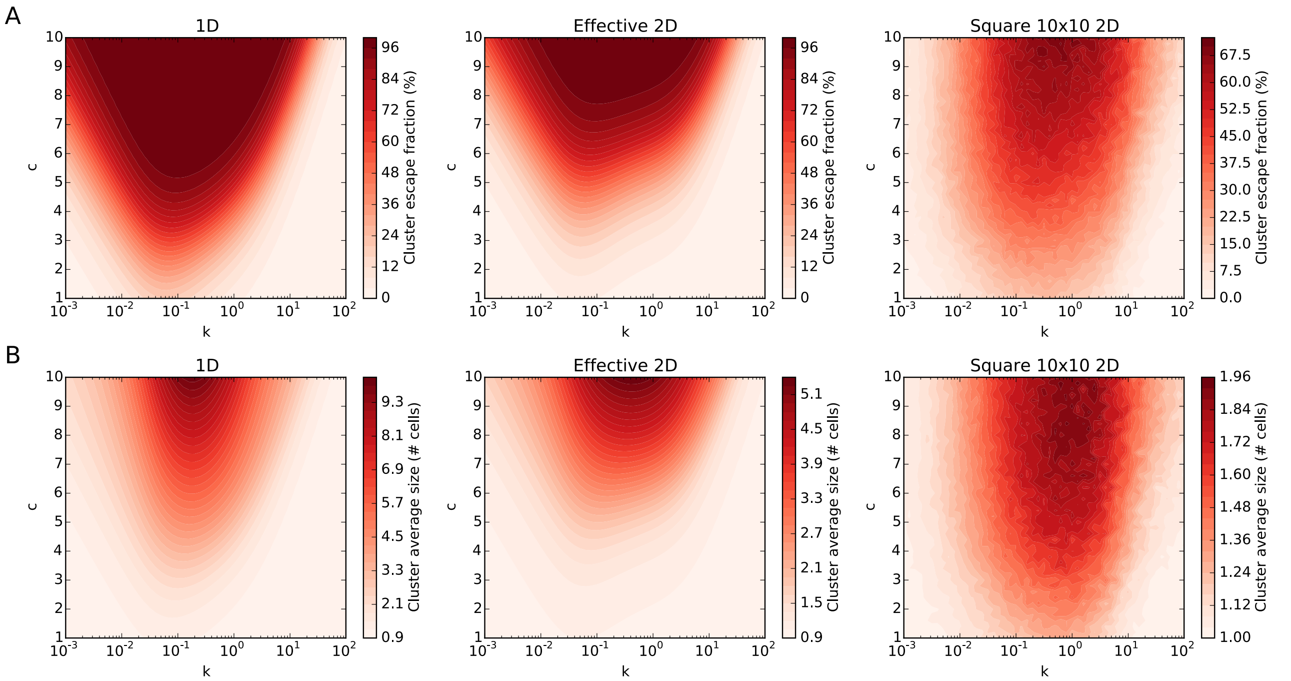


Supplementary figure 3. **Comparison between 1D, effective-2D and square-2D models. (A)** Cluster escape fraction as a function of k and c for 1D (left), effective-2D (center, same as figure 1C), and square-2D (right). **(B)** Cluster average size as a function of k and c for 1D (left), effective-2D (center, same as figure panel 2C), and square-2D (right).

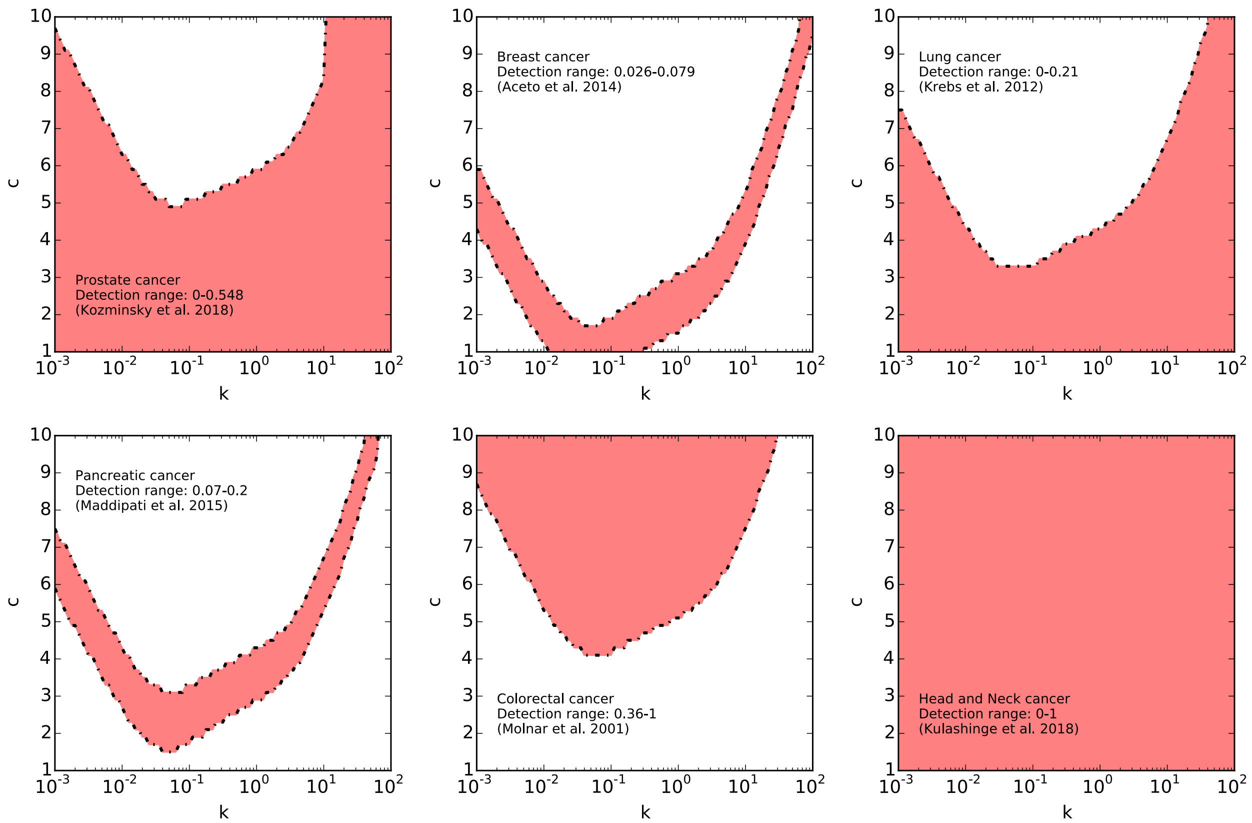


Supplementary figure 4. **CTC cluster detection ranges from independent experiments are mapped onto different regions of the parameter space.** Each panel shows the detection range of CTC clusters found in different cancers/experiments. The detection range is defined by the minimal and maximal fraction of CTC clusters observed within the cohort of patients/mouse population considered in the study. The color shading indicates the combinations of parameters in the parameter plane (k, c) of the model (Fig. 2C) for which the model exhibits a CTC cluster fraction within the experimental detection range.

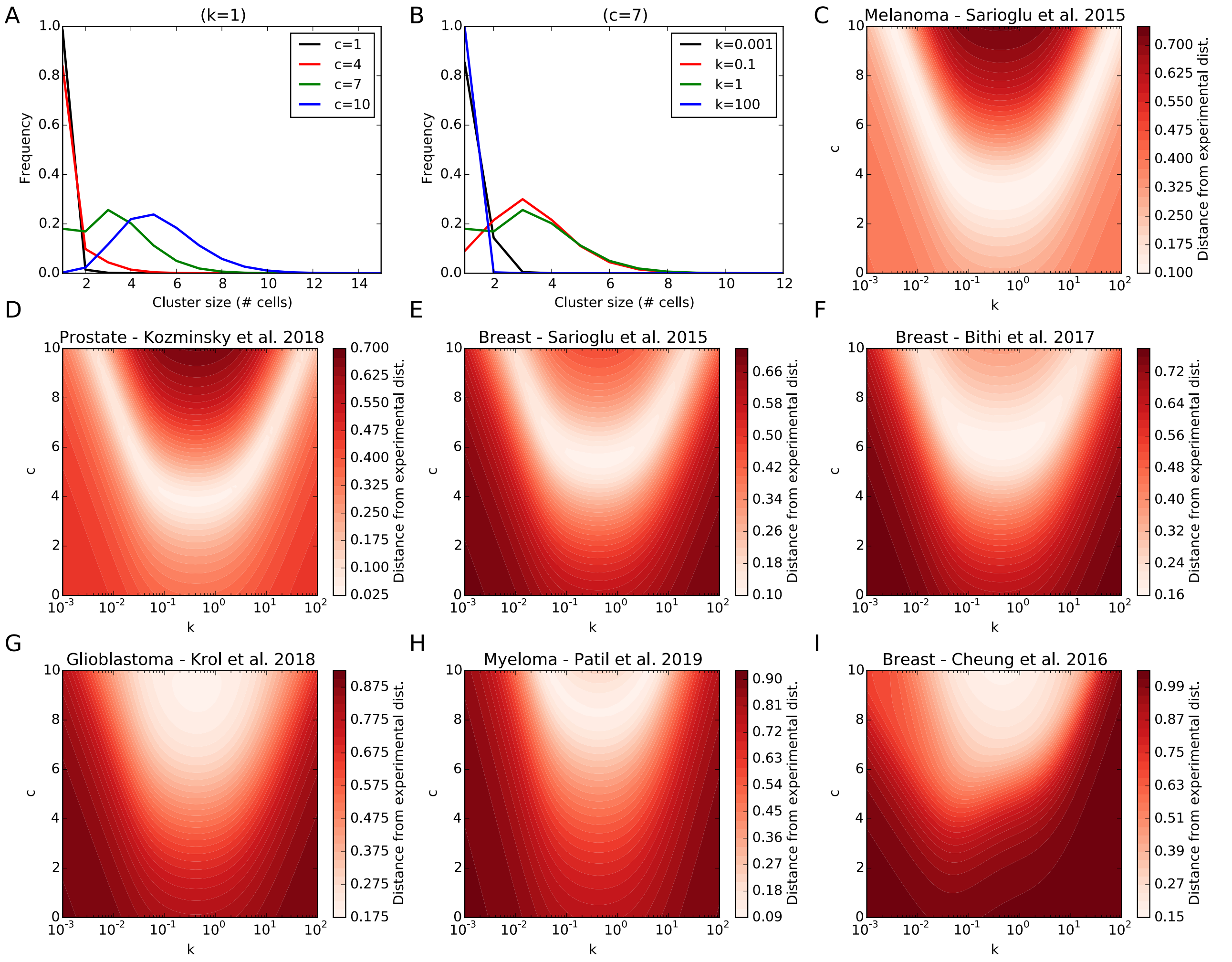


Supplementary figure 5. **Experimental cluster size distributions are mapped onto different regions of the parameter diagram. (A)** Frequency of escape as a function of cluster size for 4 values of c (1, 4, 7, 10) at a fixed EMT rate (k=1). **(B)** Frequency of escape as a function of cluster size for 4 values of k (0.01, 0.1, 1, 10) at a fixed level of cooperativity (c=1). **(C-D-E-F-G-H-I)** Distance between the experimental cluster distributions presented in Fig. 3 and the model's cluster size distribution as a function of the EMT rate (k, x-axis) and migration cooperativity (c, y-axis). The distance is defined as the square root of the square distance between the normalized experimental distribution and the model's distribution. For C-D-E-F-H, the model's distribution was renormalized by excluding the clusters of size 1 (namely, single cells) because the corresponding experiments could not detect single cells. For panel G, the renormalization excludes single cells and clusters of size n=2 because not reported in the experiment.

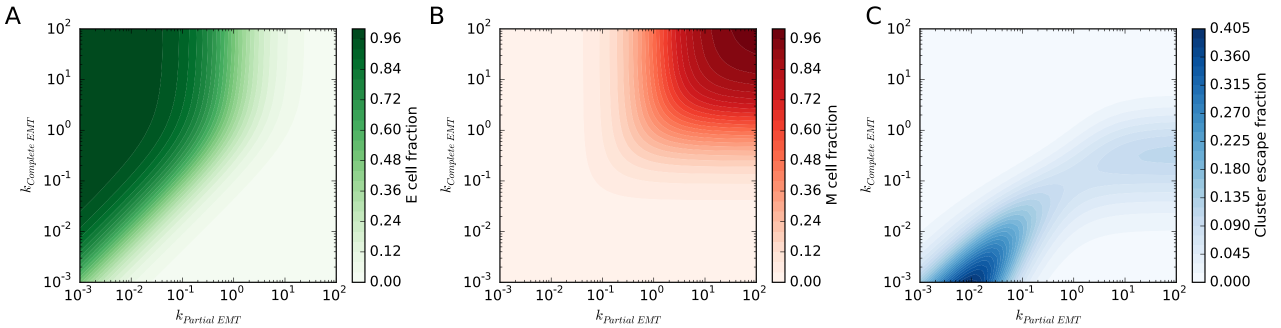


Supplementary figure 6. **Control of partial and complete EMT modulates the fraction of E, H, and M cells in the lattice.** Steady state cell fraction of epithelial cells **(A)** and mesenchymal cells **(B)** as a function of the rates of partial EMT and complete EMT in the effective-2D model with linear cooperativity (c=1). **(C)** Cluster escape as a function of the rates of partial EMT and complete EMT in the case of linear cooperativity (c=1).

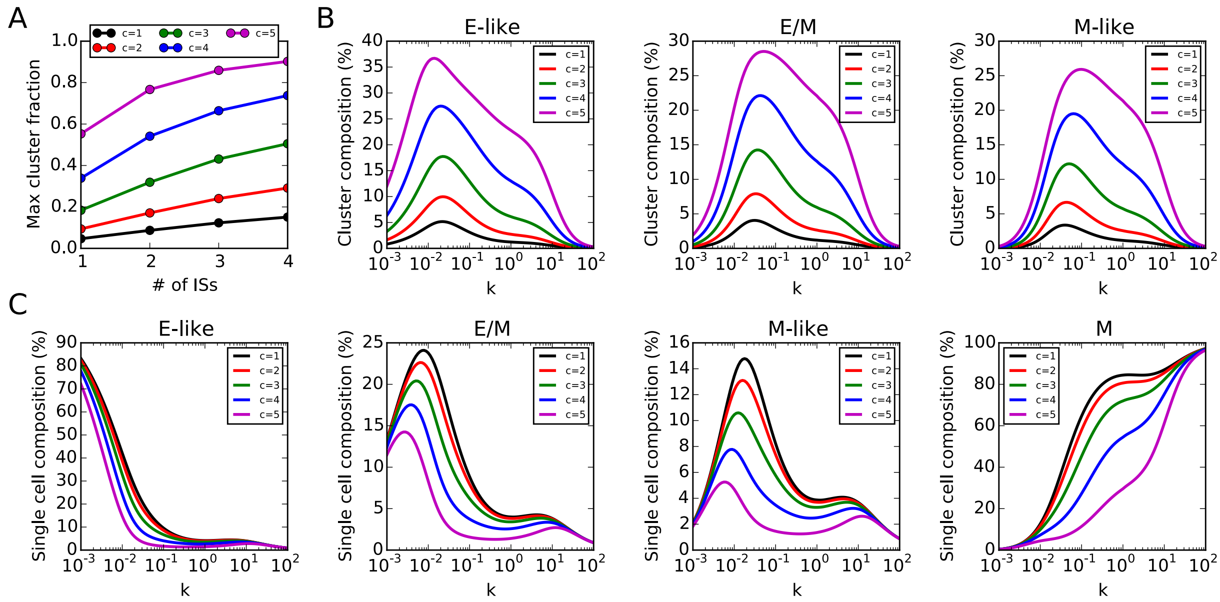


Supplementary figure 7. **The cooperative migration conserves the heterogeneous composition of clusters and single cells. (A)** Maximal cluster escape fraction as a function of the number of intermediate hybrid states (x-axis) for different levels of cooperativity (c). **(B)** Average composition of clusters as a function of the EMT rate (k) for the model with 3 intermediate hybrid states (E-like, E/M, M-like) for different levels of cooperativity (c). For a fixed value of c, the sum of the three curves equals the fraction of escape events that involve clusters. Therefore, the case c=1 (black line) represents the cluster composition shown in Figure 4C. **(C)** Fraction of single migrating cells as a function of the EMT rate (k) for the model with 3 intermediate hybrid states and different levels of cooperativity (the cell states that can migrate as single cells are E-like, E/M, M-like and M). For a fixed value of c, the sum of the four curves equals the fraction of escape events that involve single cells. Therefore, the case c=1 (black line) represents the fractions of single cells shown in Figure 4D.

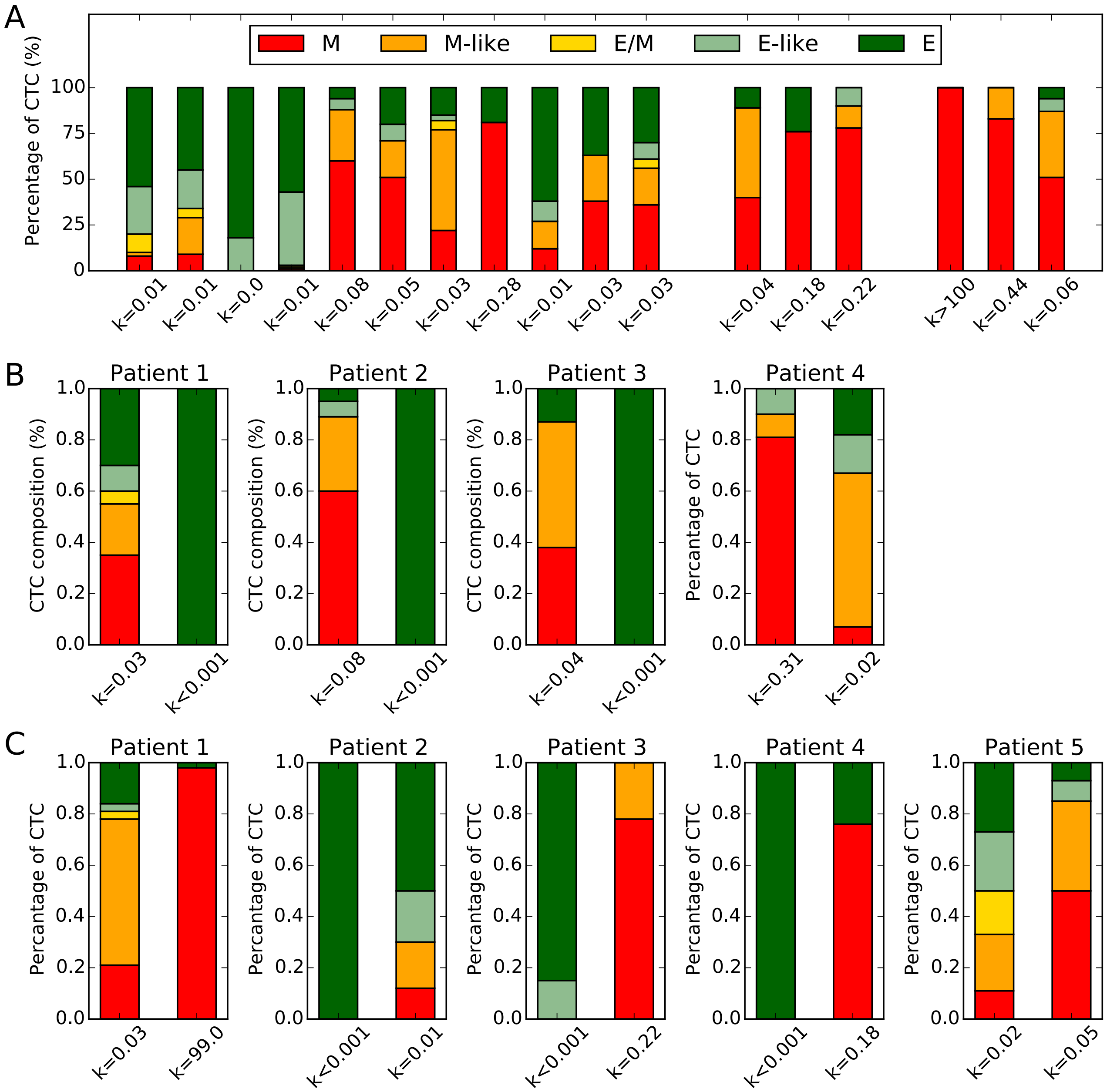


Supplementary figure 8**. CTC percentages of breast cancer patients map onto different EMT rates of the model. (A)** The percentage of epithelial, E-like, E/M, M-like and mesenchymal cells found in the bloodstream of breast cancer patients (adapted from Yu, Bardia *et al*. [19]). In the original paper, CTCs were classified as E, E>M, E=M, M>E, M. These fractions were compared to the fractions of escaping single cells in the model (such as in Fig. 4D). Since the E state is not motile in the model, we considered a case with 4 intermediate hybrid states, so that the 4 intermediates and the mesenchymal state were compared to the 5 cell fractions presented in the experiment (see Methods). The labels on the x-axis show the value of rate k that generate a fraction of escaping cell phenotypes as close as possible to the experimental data. The three groups come from different breast cancer subtypes (ER/PR1, Her2+, TN). **(B)** Percentage of CTCs pre- and post-treatment for 4 patients that exhibited positive response to treatment (adapted from Yu, Bardia *et al*. [19]). The corresponding value of k always decreases after treatment. **(C)** Percentage of CTCs pre-and post-treatment for 5 patients that exhibited negative response to treatment and underwent tumor progression instead, (adapted from Yu, Bardia *et al*. [19]). The corresponding value of k always increases after treatment.

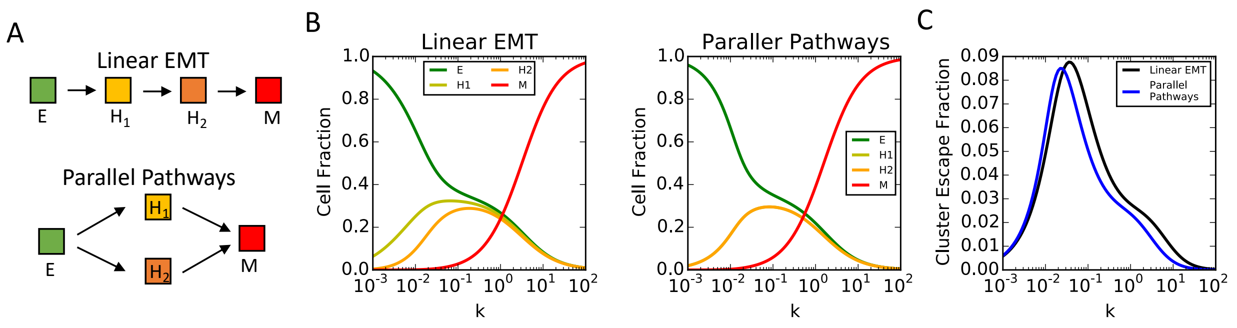


Supplementary figure 9. **Comparison between the 'linear chain' multistate EMT model and the EMT model with multiple pathways. (A)** Two intermediate hybrid states ($H_1$ and $H_2$) are considered. The states $H_1$ and $H_2$ have the same adhesion properties in the two models for a fair comparison (see supplementary section I for a discussion of the parameters used). Comparing steady state fractions of cells **(B)** in different states as well as cluster escape fraction **(C)** shows slight differences between the two models (in particular, the fraction of cells in the $H_1$ and $H_2$ states are exactly the same in the 'Parallel pathways' model). The cooperation index is c=1 in both models for this comparison.

| Panel | n=1 | n=2 | n=3 | n=4 | n=5 | n=6 | n=7 | n=8 | n=9 | ≥10 |
|---|---|---|---|---|---|---|---|---|---|---|
| 3A | 0.924 | 0.054 | 0.016 | 0.004 | >0.001 | >0.001 | >0.001 | >0.001 | >0.001 | >0.001 |
| 3B | 0.837 | 0.098 | 0.044 | 0.014 | 0.004 | 0.001 | >0.001 | >0.001 | >0.001 | >0.001 |
| 3C | 0.547 | 0.181 | 0.149 | 0.076 | 0.030 | 0.010 | 0.003 | >0.001 | >0.001 | >0.001 |
| 3D | 0.413 | 0.194 | 0.196 | 0.116 | 0.052 | 0.019 | 0.006 | 0.002 | >0.001 | >0.001 |
| 3E | 0.029 | 0.077 | 0.214 | 0.261 | 0.203 | 0.119 | 0.058 | 0.025 | 0.009 | 0.005 |
| 3F | 0.014 | 0.053 | 0.182 | 0.255 | 0.221 | 0.143 | 0.075 | 0.034 | 0.013 | 0.007 |
| 3G | 0.003 | 0.022 | 0.117 | 0.219 | 0.238 | 0.184 | 0.112 | 0.058 | 0.026 | 0.017 |

Supplementary Table 1. **Cluster escape flux percentage as a function of cluster size.** Different rows show the percentage of escape flux predicted by the model as a function of cluster size for the different curves shown in Fig. 3A-G. Since cluster size is a discrete, integer variable, the escape flux percentage of clusters of a certain size S is simply the value of the cluster size distribution at x-coordinate x=S.
